# Problematic usage of the Zhang and Luck mixture model

**DOI:** 10.1101/268961

**Authors:** Wei Ji Ma

## Abstract

A common method, due to Zhang and Luck (2008), for analyzing delayed-estimation data with a circular stimulus variable is to fit a mixture of a Von Mises distribution and a uniform distribution. The uniform distribution represents random guesses, presumably made when an item is not kept in memory. When I generate synthetic data from a variable-precision model with zero guessing, the method estimates the guess rate to be nonzero and often high. This is due to model mismatch: the fitted model is not matched to the data-generating (true) model. In real data, this could be a problem if one considers the variable-precision model a plausible candidate model and draws conclusions based on the estimated guess rates. I describe five solutions to this problem: analyzing the residual, ruling out the variable-precision model, robust inference, fitting a hybrid model, and using model-free statistics. I hope that these solutions can contribute to good data analysis practices in the study of working memory.

## Delayed estimation

Delayed estimation is a behavioral paradigm to study working memory (Blake, Cepeda, & Hiris, 1997; Prinzmetal, Amiri, Allen, & Edwards, 1998; Wilken & Ma, 2004). An observer is presented with one or more stimuli, either simultaneously or sequentially. The value of each stimulus is drawn from a (near-)continuous space. After a working memory delay, the participant reports the values of one or more of the remembered stimuli in the (near-)continuous space. Unlike categorical tasks such as change detection, delayed estimation probes the contents of working memory at high resolution: the main dependent measure is a (near-)continuous estimation error. Delayed estimation has been used to investigate, among others, the effects of set size (Wilken & Ma, 2004; Zhang & Luck, 2008), delay time (Zhang & Luck, 2009; Pertzov, Manohar, & Husain, 2017; Shin, Zou, & Ma, 2017), cueing (Zhang & Luck, 2008; P. M. Bays, 2014), reward (Zhang & Luck, 2011; Ye et al., 2017), and number of features (Fougnie & Alvarez, 2011; Swan, Collins, & Wyble, 2016; Shin & Ma, 2016) on the quality of working memory.

## The Zhang and Luck mixture model

Often, the stimulus variable in delayed estimation is chosen to be periodic: in particular, common choices are color on a color wheel, and orientation. For such variables, Zhang and Luck introduced a model to analyze the errors in a given condition (Zhang & Luck, 2008). The method starts with the assumption that on some proportion *w* of trials, the observer has no memory of the stimulus and therefore makes a random guess. These trials will produce a uniform distribution of estimation error. The model further assumes that on non-guess trials, estimation error follows a Von Mises distribution (a circular version of the Gaussian distribution). Taken together, the model’s probability density of the estimation error ϵ ∈ [0, 2π) is a mixture distribution

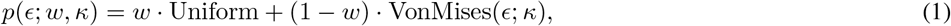

where

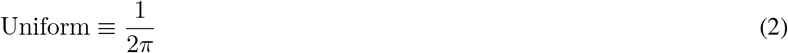

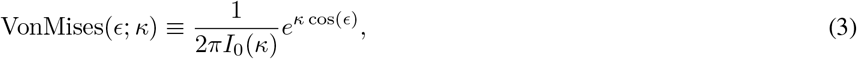

where *I_0_* is the modified Bessel function of the first kind of order 0. Free parameters are the concentration parameter *κ* and the *guess rate w*. An example of a fit is shown in Fig. 1.

Over the past 10 years, fitting the Zhang and Luck mixture model (ZL model) has become a standard way to analyze delayed-estimation data (e.g. (Rademaker, Tredway, & Tong, 2012; Suchow, Brady, Fougnie, & Alvarez, 2013; P. Bays, Catalao, & Husain, 2009; Pertzov et al., 2017; Gorgoraptis, Catalao, Bays, & Husain, 2011; Emrich, Lockhart, & Al-Aidroos, 2017; Weber, Peters, Hahn, Bledowski, & Fiebach, 2016; Zhang & Luck, 2009; Fougnie & Alvarez, 2011; T. F.Brady, Konkle, Gill, Oliva, & Alvarez, 2013; Zokaei, Gorgoraptis, Bahrami, Bays, & Husain, 2011; Swan et al., 2016; Zhang & Luck, 2011; Ye et al., 2017; Emrich, Riggall, LaRocque, & Postle, 2013)), and the claims of many studies are based on comparing the estimates of *κ* and *w* across conditions.

**Figure 1:**
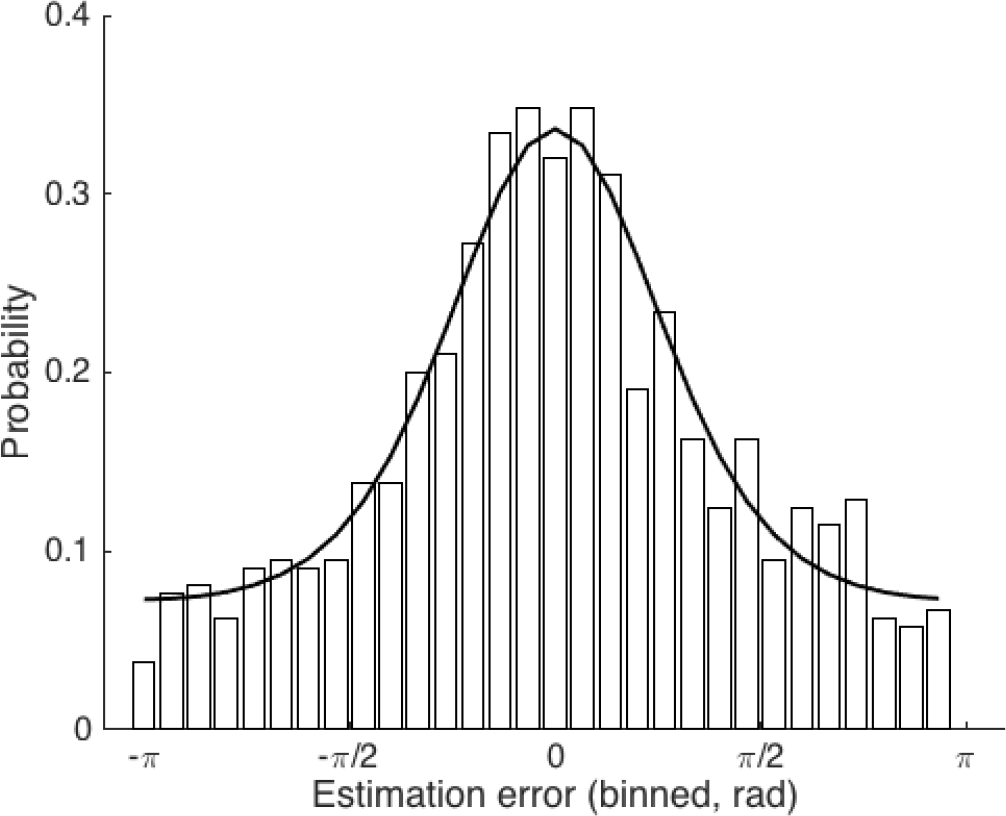
Histogram of estimation errors in a simulated delayed-estimation experiment with 1000 trials (bars), and mixture model fit (solid line). The data were generated from a variable-precision model with *J̅* =1 and *τ* =1. The mixture model estimates the guess rate w as 0.37, even though the true value was 0.

## The variable-precision model

The variable-precision (VP) model is another model for describing the distribution of estimation errors in delayed estimation (Van den Berg, Shin, Chou, George, & Ma, 2012; Fougnie, Suchow, & Alvarez, 2012). In this model, the distribution is an infinite mixture of Von Mises distributions, each with a different concentration parameter *κ*:

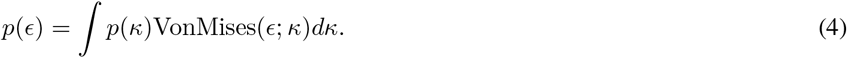

The distribution *P*(*κ*) is a probability density function on the positive real line. A natural choice for *P*(*κ*) is to assume a gamma distribution. ^1^This distribution has two parameters; one possible parametrization is in terms of a mean parameter *J̅* and a scale parameter **τ**. Many factors have been proposed to contribute to variability in *κ*, including heteroskedasticity (Pratte, Park, Rademaker, & Tong, 2017; Bae, Olkkonen, Allred, Wilson, & Flombaum, 2014), variability in spike count (P. M. Bays, 2014), variability in spike rate (Van den Berg et al., 2012), fluctuations in attention (Van den Berg et al., 2012), variability in memory decay (Fougnie et al., 2012), and contextual effects (T. Brady & Alvarez, 2015).

## Results

I simulated data from a VP model as specified above. Mean precision *J̅* was 1, 2, or 4. I varied the scale parameter *τ* linearly from 0.01 to 10, with a total of 10 values. For each combination of *J̅* and **τ**, I simulated 10 data sets. To minimize finite-sample effects, I simulated 10,000 trials per data set, many more than in a typical psychophysics experiment. On each trial, I first drew a value of *κ* from a gamma distribution with parameters *J̅* and **τ**, and then a value of ∊ from a Von Mises distribution with concentration parameter *κ*. There was no mechanism for guessing. After generating a data set, I fitted the ZL mixture model, Eq. (1), to the 10,000 trials in the data set, using maximum-likelihood estimation as implemented using Bayesian Adaptive Direct Search (Acerbi & Ma, 2017).

Figure 2 shows the estimate of the guess rate, *w*, for each data set as well as averaged across the 10 data sets with the same parameters. Across many parameter combinations, the guess rate is overestimated relative to the true value of 0. Overestimation was more severe when *J̅* was lower or **τ** was higher. This makes sense, because lower *J̅* and higher **τ** tend to produce more low-precision (low *κ*) representations, and the ZL model mistakes low-precision representations for zero-precision representations (Van den Berg et al., 2012; Van den Berg, Awh, & Ma, 2014). As a consequence, if the estimated guess rate were to increase between two conditions, this could be due to a decrease in *J̅* or an increase in **τ**, instead of due to an increase in the true guessing rate. Put more starkly, if I had included true guessing in the data generation, I would have found that the estimated guess rate could increase between two conditions even when the true guess decreased (in combination with a decrease in *J̅* or an increase in **τ**). Thus, if errors follow the variable-precision model, the guess rate estimated in the ZL model is not a reliable measure of either the value of or the trend in the true guess rate. This is not necessarily a problem in itself; however, it could be if the narrative of the study relies on the estimated guess rates.

## Discussion

When fitting the ZL model to data generated from the VP model, I found severe and variable overestimation of the guess rate. A similar overestimate was previously found in a change detection task, where it was quantified using an “apparent guessing rate” (Keshvari, Van den Berg, & Ma, 2013). Fig. 2 by no means speaks to the presence or absence of true guessing (for an interesting new take on that question, see (Adam, Vogel, & Awh, 2017)); it only illustrates a pitfall in interpreting the guess rate parameter in the ZL model.

The overestimation is a straightforward consequence of *model mismatch* or *model misspecification* (Ramsey, 1969): there is a mismatch between the data-generating model (VP) and the fitted model (ZL). Model mismatch often leads to biases in parameter estimation, and such biases are by no means unique to the ZL model. For example, when psychometric curve data are generated in the presence of a lapse rate, but a model without a lapse rate is fitted, the slope of the psychometric curve is systematically underestimated (Wichmann & Hill, 2001). Similarly, if I had fitted the VP model to data generated from the ZL model, I could easily have obtained meaningless estimates.

Despite this symmetry in principle between the VP model and the ZL model, the practical usage of both models has differed greatly. The VP model is usually compared in great detail to alternative models (e.g. (Van den Berg et al., 2012, 2014; Nosofsky & Donkin, 2016; Van den Berg, Yoo, & Ma, 2017; Emrich et al., 2017; Devkar, Wright, & Ma, 2015; Keshvari et al., 2013; Pratte et al., 2017)). Moreover, in these studies, the estimates of the parameters in the variable-precision model, even if it outperforms other models, are not typically emphasized or analyzed further. By contrast, when the ZL model is fitted, it is often the only model being considered. Moreover, its parameter estimates are then used as the starting point for further analysis, such as a comparison across conditions. This way of using the ZL model seems flawed: the parameters of a model, in particular the guess rate in the ZL mixture model, can only be trusted to the extent that one believes there to be no model mismatch.

**Figure 2:**
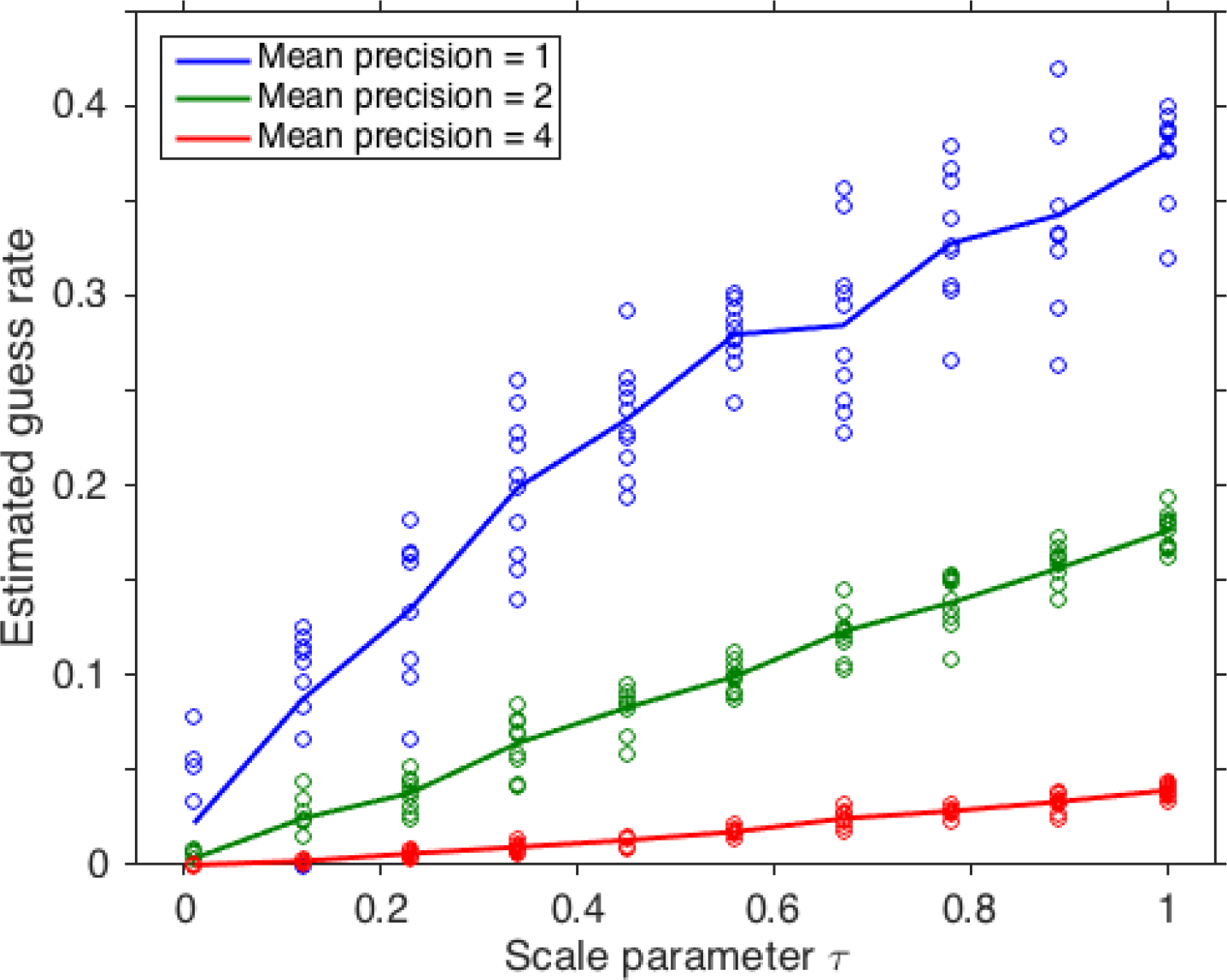
The Zhang and Luck mixture model overestimates the guess rate if the true model is a variable-precision model. The data were generated from a variable-precision model with mean precision *J̅* equal to 1, 2, or 4, and a variable scale parameter **τ**. For each parameter combination, I performed 10 simulations. Each simulated data set consisted of 10,000 trials. In every case, the true guess rate was 0.

I suggest five potential solutions to this problem:

1. Acknowledge potential model mismatch and treat the residual of the ZL model as another summary statistic (Van den Berg et al., 2012, 2014). This could complicate hypothesis testing, since this summary statistic is high-dimensional. Moreover, recognizing that the residual is structured makes the ZL model less relevant as a model of the underlying cognitive process.
2. Formally compare the ZL model against the VP model. If the ZL model wins convincingly, there is a basis for trusting and further analyzing its parameter estimates. Such a convincing win might be rare, though: usually, the VP model fits as well or somewhat better (Van den Berg et al., 2014, 2017).
3. Ensure that any inferences drawn from a comparison across conditions are independent of the fitted model (Shin et al., 2017). The disadvantages of this approach are that it is more work and that a decision must be made on how to aggregate results that are not completely consistent across models.
4. Fit a hybrid model that contains both the VP model and the ZL model as special cases. An example of such a model is a VP model with a guessing component (Van den Berg et al., 2014, 2017; Devkar et al., 2015). Indeed, this model sometimes performs better than a pure VP model, providing evidence for the presence of true guessing. Of course, this approach does not exclude the possibility of a mismatch between the hybrid model and the true model; however, it is a good way to not have to adjudicate between the ZL and VP models.
5. Forego modeling altogether and compare across conditions only model-free summary statistics, such as mean absolute error, circular variance, or circular standard deviation (Van den Berg et al., 2017; Shin et al., 2017; Rademaker et al., 2012; Pertzov et al., 2017; Emrich et al., 2017). This is my best recommendation if the purpose of a study is not model comparison but a characterization of some independent variable on working memory performance.

## Footnotes

1 An alternative is to first define precision as 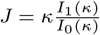, where *I*_1_ is the modified Bessel function of the first kind of order 1, and then assume a gamma distribution over *J* (Van den Berg et al., 2012). For the purpose of the present paper, this is an unnecessary complication.

## References

Acerbi, L., & Ma, W. J. (2017). Practical bayesian optimization for model fitting with bayesian adaptive direct search. eprint arXiv:1705.04405.

Adam, K. C., Vogel, E. K., & Awh, E. 2017. Clear evidence for item limits in visual working memory. Cognitive psychology, 97, 79–97.

Bae, G.-Y., Olkkonen, M., Allred, S. R., Wilson, C., & Flombaum, J. I. 2014. Stimulus-specific variability in color working memory with delayed estimation. Journal of Vision, 14(4), 7–7.

Bays, P., Catalao, R., & Husain, M. 2009. The precision of visual working memory is set by allocation of a shared resource. Journal of vision, 9(10), 7–7.

Bays, P. M. 2014. Noise in neural populations accounts for errors in working memory. Journal of Neuroscience, 34(10), 3632–3645.

van den Berg, R., Awh, E., & Ma, W. J. 2014. Factorial comparison of working memory models. Psychological review, 121(1), 124.

van den Berg, R., Shin, H., Chou, W.-C., George, R., & Ma, W. J. 2012. Variability in encoding precision accounts for visual short-term memory limitations. Proceedings of the National Academy of Sciences, 109(22), 8780–8785.

van den Berg, R., Yoo, A. H., & Ma, W. J. 2017. Fechner?s law in metacognition: A quantitative model of visual working memory confidence. Psychological review, 124(2), 197.

Blake, R., Cepeda, N. J., & Hiris, E. 1997. Memory for visual motion. Journal of Experimental Psychology: Human Perception and Performance, 23(2), 353–369.

Brady, T., & Alvarez, G. A. 2015. Contextual effects in visual working memory reveal hierarchically structured memory representations. Journal of vision, 15(15), 6–6.

Brady, T. F., Konkle, T., Gill, J., Oliva, A., & Alvarez, G. A. 2013. Visual long-term memory has the same limit on fidelity as visual working memory. Psychological Science, 24(6), 981–990.

Devkar, D. T., Wright, A. A., & Ma, W. J. 2015. The same type of visual working memory limitations in humans and monkeys. Journal of vision, 15(16), 13–13.

Emrich, S. M., Lockhart, H. A., & Al-Aidroos, N. 2017. Attention mediates the flexible allocation of visual working memory resources. Journal of Experimental Psychology: Human Perception and Performance, 43(7), 1454.

Emrich, S. M., Riggall, A. C., LaRocque, J. J., & Postle, B. R. 2013. Distributed patterns of activity in sensory cortex reflect the precision of multiple items maintained in visual short-term memory. Journal of Neuroscience, 33(15), 6516–6523.

Fougnie, D., & Alvarez, G. A. 2011. Object features fail independently in visual working memory: Evidence for a probabilistic feature-store model. Journal of vision, 11(12), 3–3.

Fougnie, D., Suchow, J. W., & Alvarez, G. A. 2012. Variability in the quality of visual working memory. Nature communications, 3, 1229.

Gorgoraptis, N., Catalao, R., Bays, P., & Husain, M. (2011). Dynamic updating of working memory resources for visual objects. Journal of Neuroscience, 31(23), 8502–8511.

Keshvari, S., van den Berg, R., & Ma, W. J. 2013. No evidence for an item limit in change detection. PLoS computational biology, 9(2), e1002927.

Nosofsky, R. M., & Donkin, C. 2016. Qualitative contrast between knowledge-limited mixed-state and variable-resources models of visual change detection. Journal of Experimental Psychology: Learning, Memory, and Cognition, 42(10), 1507.

Pertzov, Y., Manohar, S., & Husain, M. 2017. Rapid forgetting results from competition over time between items in visual working memory. Journal of Experimental Psychology: Learning, Memory, and Cognition, 43(4), 528.

Pratte, M. S., Park, Y. E., Rademaker, R. L., & Tong, F. 2017. Accounting for stimulus-specific variation in precision reveals a discrete capacity limit in visual working memory. Journal of Experimental Psychology: Human Perception and Performance, 43(1), 6.

Prinzmetal, W., Amiri, H., Allen, K., & Edwards, T. 1998. Phenomenology of attention: I. color, location, orientation, and spatial frequency. Journal of Experimental Psychology: Human Perception and Performance, 24(1), 261–282.

Rademaker, R., Tredway, C., & Tong, F. 2012. Introspective judgments predict the precision and likelihood of successful maintenance of visual working memory. Journal of Vision, 12(13), 21–21.

Ramsey, J. B. 1969. Tests for specification errors in classical linear least-squares regression analysis. Journal of the Royal Statistical Society. Series B (Methodological), 350–371.

Shin, H., & Ma, W. J. 2016. Crowdsourced single-trial probes of visual working memory for irrelevant features. Journal of vision, 16(5), 10–10.

Shin, H., Zou, Q., & Ma, W. J. 2017. The effects of delay duration on visual working memory for orientation. Journal of vision, 17(14), 10–10.

Suchow, J. W., Brady, T. F., Fougnie, D., & Alvarez, G. A. 2013. Modeling visual working memory with the memtoolbox. Journal of vision, 13(10), 9–9.

Swan, G., Collins, J., & Wyble, B. 2016. Memory for a single object has differently variable precisions for relevant and irrelevant features. Journal of vision, 16(3), 32–32.

Weber, E. M. G., Peters, B., Hahn, T., Bledowski, C., & Fiebach, C. J. 2016. Superior intraparietal sulcus controls the variability of visual working memory precision. Journal of Neuroscience, 36(20), 5623–5635.

Wichmann, F. A., & Hill, N. J. 2001. The psychometric function: I. fitting, sampling, and goodness of fit. Perception & psychophysics, 63(8), 1293–1313.

Wilken, P., & Ma, W. J. 2004. A detection theory account of change detection. Journal of Vision, 4(12), 11.

Ye, C., Hu, Z., Li, H., Ristaniemi, T., Liu, Q., & Liu, T. 2017. A two-phase model of resource allocation in visual working memory. Journal of Experimental Psychology: Learning, Memory, and Cognition, 43(10), 1557.

Zhang, W., & Luck, S. J. 2008. Discrete fixed-resolution representations in visual working memory. Nature, 453(7192), 233.

Zhang, W., & Luck, S. J. 2009. Sudden death and gradual decay in visual working memory. Psychological science, 20(4), 423–428.

Zhang, W., & Luck, S. J. 2011. The number and quality of representations in working memory. Psychological Science, 22(11), 1434–1441.

Zokaei, N., Gorgoraptis, N., Bahrami, B., Bays, P. M., & Husain, M. 2011. Precision of working memory for visual motion sequences and transparent motion surfaces. Journal of vision, 11(14), 2–2.

